# Shifting mutational constraints in the SARS-CoV-2 receptor-binding domain during viral evolution

**DOI:** 10.1101/2022.02.24.481899

**Authors:** Tyler N. Starr, Allison J. Greaney, William W. Hannon, Andrea N. Loes, Kevin Hauser, Josh R. Dillen, Elena Ferri, Ariana Ghez Farrell, Bernadeta Dadonaite, Matthew McCallum, Kenneth A. Matreyek, Davide Corti, David Veesler, Gyorgy Snell, Jesse D. Bloom

## Abstract

SARS-CoV-2 has evolved variants with substitutions in the spike receptor-binding domain (RBD) that impact its affinity for ACE2 receptor and recognition by antibodies. These substitutions could also shape future evolution by modulating the effects of mutations at other sites—a phenomenon called epistasis. To investigate this possibility, we performed deep mutational scans to measure the effects on ACE2 binding of all single amino-acid mutations in the Wuhan-Hu-1, Alpha, Beta, Delta, and Eta variant RBDs. Some substitutions, most prominently N501Y, cause epistatic shifts in the effects of mutations at other sites, thereby shaping subsequent evolutionary change. These epistatic shifts occur despite high conservation of the overall RBD structure. Our data shed light on RBD sequence-function relationships and facilitate interpretation of ongoing SARS-CoV-2 evolution.

## Main text

The SARS-CoV-2 spike receptor-binding domain (RBD) has undergone rapid evolution since the virus’s emergence (*1*). We previously used deep mutational scanning to experimentally measure the impact of all single amino-acid mutations on the ACE2-binding affinity of the ancestral Wuhan-Hu-1 RBD (*2*). These measurements have helped inform surveillance of SARS-CoV-2 evolution. For example, we identified the N501Y mutation as enhancing ACE2-binding affinity prior to the emergence of this consequential mutation in the Alpha variant (*3*).

However, as proteins evolve, the impacts of individual amino acid mutations can shift due to epistasis (*4*). For example, the same N501Y mutation that enhances SARS-CoV-2 binding to ACE2 severely impairs ACE2 binding by SARS-CoV-1 and other divergent sarbecoviruses (*5*). Furthermore, N501Y epistatically enabled other affinity-enhancing mutations that emerged in the Omicron variant of SARS-CoV-2 (*6*–*8*). To more systematically understand how epistasis shifts the effects of mutations, we performed deep mutational scans to measure the impacts of all individual amino-acid mutations in SARS-CoV-2 variant RBDs.

We constructed comprehensive site-saturation mutagenesis libraries in the ancestral Wuhan-Hu-1 RBD and RBDs from four variants: Alpha (N501Y), Beta (K417N+E484K+N501Y), Delta (L452R+T478K) and Eta (E484K). We cloned these mutant libraries into a yeast-surface display platform and determined the impact of every amino acid mutation on ACE2-binding affinity and yeast surface-expression levels via FACS and high-throughput sequencing (Figs. S1, S2 and Data S1) (*2*). The effect of each mutation on ACE2 binding is shown in Fig. 1, and an interactive version of this figure is available at https://jbloomlab.github.io/SARS-CoV-2-RBD_DMS_variants/RBD-heatmaps/. We used monomeric ACE2 ectodomain to measure 1:1 binding affinities, which provide more granularity to reveal affinity-enhancing effects compared to our previous measurements using the natively dimeric ACE2 ligand (Fig. S1F) (*2*).

**Fig. 1.**
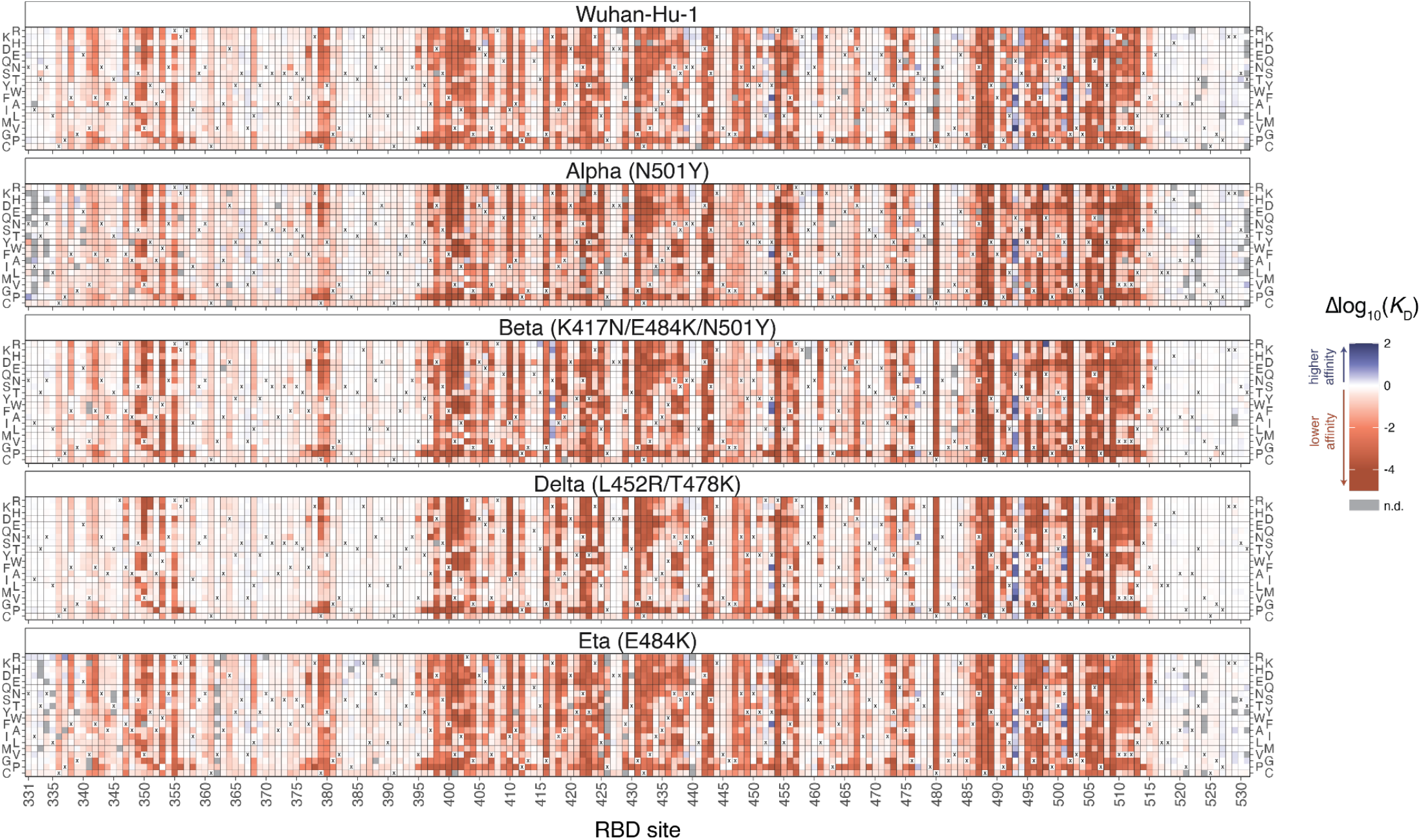
Deep mutational scanning maps of ACE2-binding affinity for all single amino acid mutations in five SARS-CoV-2 RBD variants. The impact on ACE2 receptor-binding affinity (Δlog_10_(*K*_D_)) of every single amino-acid mutation in SARS-CoV-2 RBDs, as determined by high-throughput titration assays (Fig. S1). The wildtype amino acid in each variant is indicated with an “x”, and gray squares indicate missing mutations in each library. An interactive version of this map is at https://jbloomlab.github.io/SARS-CoV-2-RBD_DMS_variants/RBD-heatmaps/, and raw data are in Data S1. The effects of mutations on RBD surface expression are in Fig. S2.

We identified sites where the impacts of mutations differ between RBD variants (Fig. 2 and Figs. S3, S4). Epistatic shifts in mutational effects on ACE2 binding are primarily attributable to the N501Y mutation: the effects of mutations in the Delta (L452R+T478K) and Eta (E484K) RBDs are similar to those in the ancestral Wuhan-Hu-1 RBD, and the differences in the Beta (K417N+E484K+N501Y) RBD are largely recapitulated in the Alpha RBD containing N501Y alone (Fig. 2A,B). One exception is a unique epistatic shift in the effects of mutations to serine or threonine at site 419 in the Beta RBD that introduce an N-linked glycosylation sequon when an asparagine is present via the K417N mutation (Fig. S3D).

**Fig. 2.**
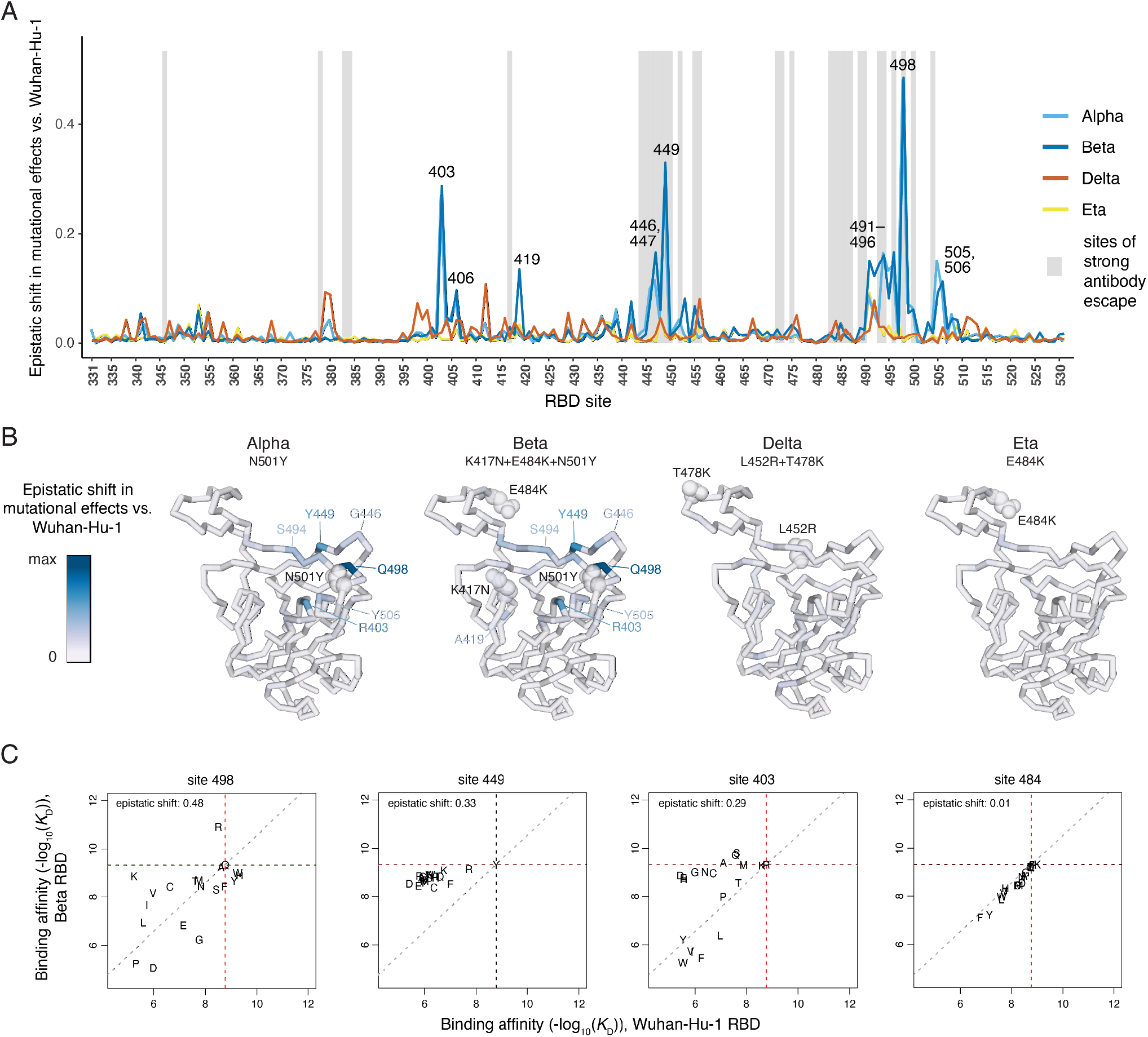
Epistatic shifts in mutational effects across RBD variants. (**A**) The shift in mutational effects on ACE2 binding at each RBD site between the indicated variant and Wuhan-Hu-1. An interactive version of this plot is at https://jbloomlab.github.io/SARS-CoV-2-RBD_DMS_variants/epistatic-shifts/. The epistatic shift is calculated as the Jensen-Shannon divergence in the set of Boltzmann-weighted affinities for all amino acids at each site. Gray shading indicates sites of strong antibody escape based on prior deep mutational scanning of the Wuhan-Hu-1 RBD (*9*). (**B**) Ribbon diagram of the Wuhan-Hu-1 RBD structure (PDB 6M0J) colored according to epistatic shifts. Labeled spheres indicate residues that are mutated in each RBD variant. (**C**) Mutation-level plots of epistatic shifts at sites of interest. Each scatter plot shows the measured affinity of all 20 amino acids in the Beta versus Wuhan-Hu-1 RBD. Red dashed lines mark the parental RBD affinities, and the gray dashed line indicates the additive (non-epistatic) expectation. Epistatic shifts can reflect idiosyncratic mutation-specific shifts (e.g., site 498) or global changes in mutational sensitivity at a site (e.g., site 449). Site 484 does not have a substantial epistatic shift and is shown for comparison. See Fig. S3 for scatterplots of additional sites of interest. See Fig. S4 for epistatic shifts in mutational effects on RBD expression.

The RBD sites that exhibit notable epistatic shifts due to N501Y fall into three structural groups (Fig. 2B). The largest shift in mutational effects is at the direct N501-contact residue Q498 (Fig. 2C), together with further epistatic shifts at sites 491-496 comprising the central beta strand of the ACE2-contact surface (Fig. 2B and Fig. S3A). A second cluster of sites exhibiting epistatic shifts in the presence of N501Y include 446, 447, and 449, which do not directly contact N501 but are spatially adjacent to residue 498 (Fig 2B,C and Fig. S3B). A third group of sites that epistatically shift due to N501Y includes residue R403 (Fig. 2C), together with several residues (505, 506, and 406) that form an interaction network linking site 501 to site 403 (Fig. S3C).

Some of these epistatic shifts are of clear relevance during the recent evolution of SARS-CoV-2. One of the strongest epistatic shifts is the potentiation of Q498R by N501Y (Figs. 2C, 3A). Although Q498R alone is mildly deleterious for ACE2 affinity in the Wuhan-Hu-1 RBD, it confers a 25-fold enhancement in affinity when present in conjunction with N501Y (which itself improves binding 15-fold), such that the double mutant has a 387-fold increased binding affinity. The Q498R/N501Y double mutation was first discovered in directed evolution studies (*6*) and is present in the RBD of the Omicron BA.1 and BA.2 variants (*8*). The epistasis between these two mutations is crucial for enabling the Omicron RBD to bind ACE2 with high affinity despite having a large number of mutations (*10*–*12*). Specifically, the set of mutations in the Omicron RBD are predicted to strongly impair ACE2 affinity based on their summed single-mutant effects in Wuhan-Hu-1 (Fig. 3B, left panel), but their summed single-mutant effects in the Beta background (which has N501Y) is about zero, consistent with the actual affinity of the Omicron RBD for ACE2 (Fig. 3B, right panel). Therefore, the affinity buffer conferred by the epistatic Q498R/N501Y pair enables the Omicron spike to tolerate other mutations that are deleterious for ACE2 binding (Fig. 3B and Fig. S5A) but contribute to antibody escape (Fig. S5B,C) (*9*). Consistent with these affinity measurements, introducing R498Q and Y501N reversions into the Omicron BA.1 spike reduces cell entry by spike-pseudotyped lentiviral particles, demonstrating that the remaining Omicron RBD mutations are deleterious without buffering by Q498R/N501Y (Fig. 3C and Fig. S6A,B).

**Fig. 3.**
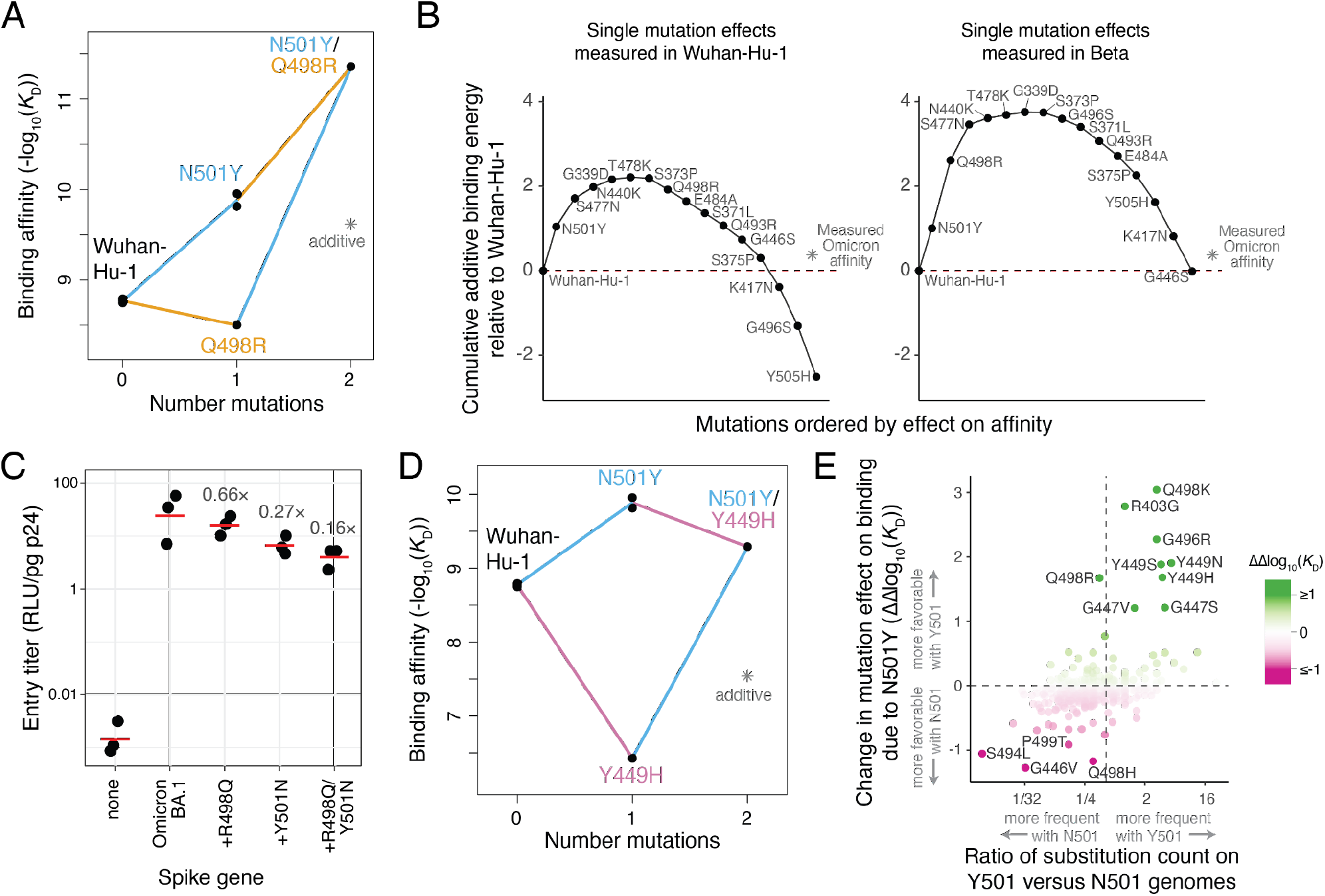
Functional and evolutionary relevance of epistatic interactions. (**A**) Double mutant cycle diagram illustrating the favorable interaction (positive sign epistasis) between N501Y and Q498R. Asterisk indicates expected double-mutant binding affinity assuming additivity. (**B**) Affinity-buffering of Omicron BA.1 mutations. Each diagram shows the cumulative addition of individually measured effects on ACE2-binding affinity (Δlog_10_(*K*_D_)) for each single RBD substitution in Omicron BA.1 as measured in the Wuhan-Hu-1 (left) or Beta (right) RBDs. Red line marks the Wuhan-Hu-1 affinity, and asterisk the actual affinity of the Omicron BA.1 RBD relative to Wuhan-Hu-1 as measured in (*10*). See also Fig. S5A–C. (**C**) Efficiency of entry of Omicron BA.1 (or reversion mutant) spike-pseudotyped lentivirus on a HEK-293T cell line expressing low levels of ACE2 (Fig. S6A,B). Labels indicate fold-decrease in geometric mean (red bar) of biological triplicate measurements. (**D**) Double mutant cycle illustrating positive epistasis between N501Y and Y449H. (**E**) Impact of epistasis on SARS-CoV-2 sequence evolution. Plot illustrates the change in a mutation’s effect between Alpha (N501Y) versus Wuhan-Hu-1 deep mutational scanning data, versus the ratio in number of observed occurrences of the substitution in genomes containing N501 versus Y501 in a global SARS-CoV-2 phylogeny as of 7 February, 2022 (*20*). A pseudocount was added to all substitution counts to enable ratio comparisons, and substitutions that were observed <2 times in total are excluded. Color scale reinforces the ΔΔlog_10_(*K*_D_) metric on the y-axis. Labeled mutations are those with |ΔΔlog_10_(*K*_D_)| > 0.9. Vertical line at *x* ~ 0.5 marks equal relative occurrence on Y501 versus N501 genomes given the larger number of substitutions that have been observed on N501 genomes to date.

There is also evolutionary relevance of the epistasis of N501Y with mutations on the 446-449 loop, which comprises the epitope for an important class of human antibodies (*13*, *14*). Although mutations to G446 escape this class of antibodies in the Wuhan-Hu-1 RBD (*14*, *15*), these mutations incur a strong ACE2-binding deficit in the N501Y background (Figs. S3B and S5D). Conversely, mutations to Y449 are largely deleterious in the Wuhan-Hu-1 RBD but are better tolerated when accompanied by N501Y (Figs. 2C, 3D). Mutations to Y449 can escape monoclonal antibodies (Fig. S6C–E) (*13*, *16*) and reduce neutralization by polyclonal sera (*17*, *18*). Mutations to Y449 have been described in several variants that also contain N501Y, including the C.1.2, A.29, and B.1.640 lineages (*17*, *19*).

To more systematically examine how epistatic shifts impact patterns of sequence variation during SARS-CoV-2 evolution, we counted the occurrence of substitutions on a global SARS-CoV-2 phylogeny (*20*). We observed that mutations more often occurred in backgrounds containing the amino acid at site 501 with which they had more favorable epistasis (Fig. 3E). Therefore, our data enables identification of mutations like those at site Y449 whose evolutionary relevance may grow if N501Y variants continue to predominate.

However, other common combinations of mutations are not involved in specific epistatic interactions with respect to ACE2 affinity. For instance, substitutions at sites 417, 484, and 501 arose together in the Beta and Gamma variants. Early studies conflicted on whether there is epistasis among these mutations with respect to ACE2 binding (*6*, *21*, *22*). Our data demonstrate strict additivity among mutations at these three sites with respect to ACE2 binding (Figs. S3E and S5E). The co-occurrence of mutations at these three sites in SARS-CoV-2 variants may instead reflect antigenic selection for E484 and K417 (which escape different classes of neutralizing antibodies (*13*), while N501Y might globally compensate for the affinity-decreasing effect of K417 mutations.

To examine the structural basis for epistatic shifts in mutational effects, we determined a crystal structure of the ACE2-bound Beta RBD (plus antibodies S304 and S309) at 2.45Å resolution (Table S1) and compared it to previously determined ACE2-bound Wuhan-Hu-1 and Beta structures (*23*, *24*). These comparisons do not reveal clear structural perturbations that explain epistatic shifts between the Wuhan-Hu-1 and Beta RBDs (Fig. S7). The K417N, E484K and N501Y substitutions cause minimal changes to the Beta RBD backbone (Fig. 4A and Fig. S8A). Furthermore, we did not detect any correlation between structural displacement of backbone or sidechain atoms in variant RBD structures and epistatic shifts in mutational effects (Fig. 4B and Fig. S8B–E), indicating that epistatic shifts occur despite conservation of the global and local RBD structure.

**Fig. 4.**
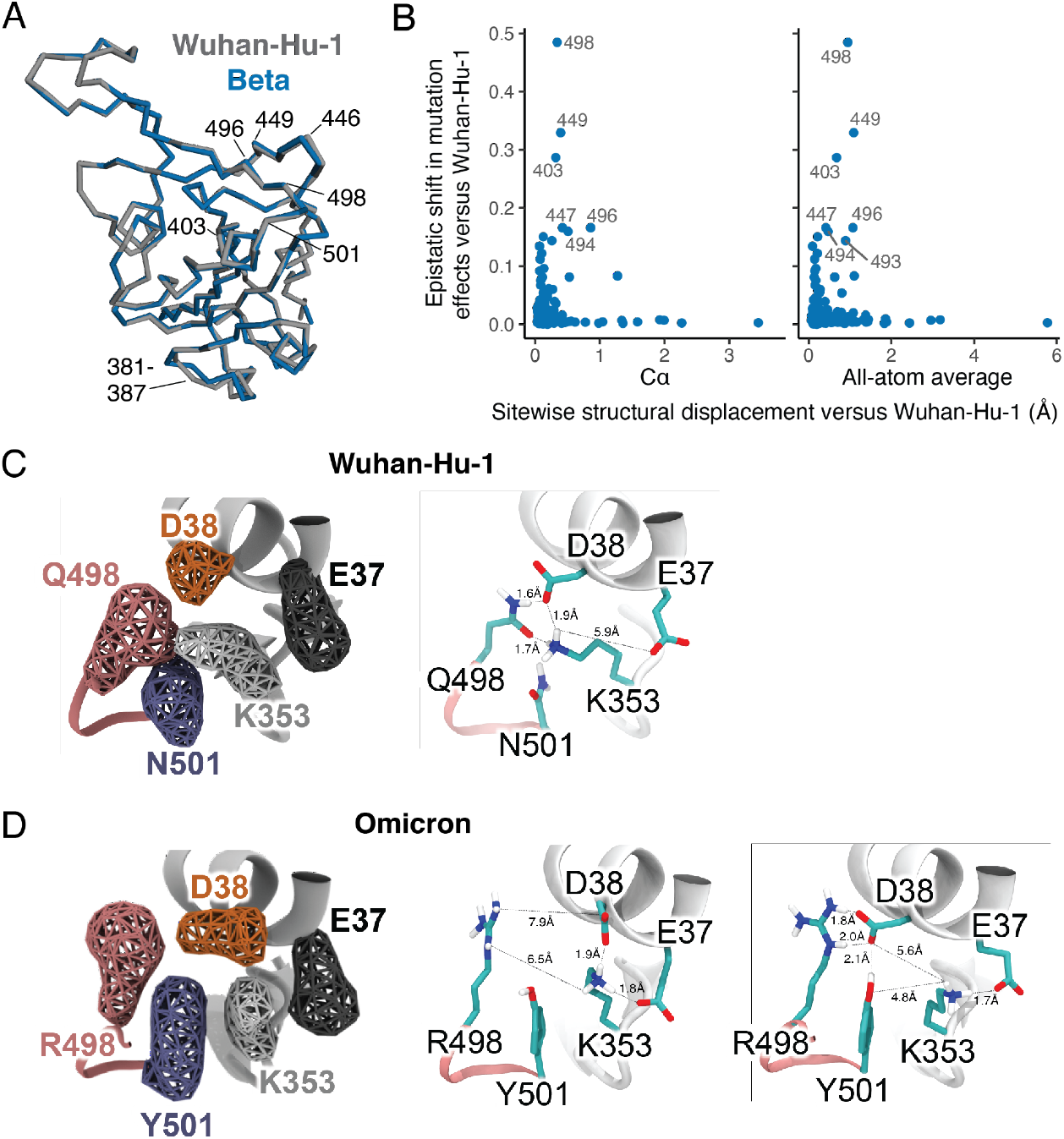
Epistatic shifts are not accompanied by large structural perturbations. (**A**) Global alignment of the Wuhan-Hu-1 (PDB 6M0J) and Beta (PDB 7EKG) RBD backbones. Key sites are labeled. (**B**) Correlation between the extent of epistatic shift in mutational effects at a site and its structural perturbation in Beta versus Wuhan-Hu-1 RBDs (backbone Cα or all-atom average displacement from aligned X-ray crystal structures). See Figs. S7, S8 for additional details. (**C**, **D**) Details from molecular dynamics simulation of Wuhan-Hu-1 (C) or Omicron (D) RBD bound to ACE2. Volumetric maps (left) show the 3D space occupied by key residues over the course of simulation. Representative snapshots (right) illustrate a dominant polar contact network in Wuhan-Hu-1 between residues K353, D38, and Q498 (C) is disrupted in Omicron, where D38 can either salt bridge K353 or R498, but not both simultaneously (D).

To explore the cause of epistasis between Q498R and N501Y (Fig. 3A), we examined structures of Wuhan-Hu-1 and Omicron RBDs bound to ACE2 (*12*, *23*) complemented by molecular dynamics simulation (Fig. 4C,D and Fig. S9). The Wuhan-Hu-1 structure features a stable polar contact network between ACE2 residues D38 and K353 and RBD residue Q498 (Fig. 4C). The N501Y substitution present in Omicron disrupts this polar contact network, as the larger Y501 packs with the aliphatic base of R498 and K353, sterically blocking K353 from forming the contact bridging D38 and RBD residue 498 (Fig. 4D). In the Omicron structure, D38 instead alternates salt bridge contacts with K353 and the substituted R498 side chain (Fig. 4D). N501Y may therefore potentiate the affinity enhancement of Q498R by improving the energetic benefit of the stronger R498:D38 salt bridge when D38 can no longer jointly coordinate K353 in this conformation. Further support of this mechanism can be seen in the similar pattern of N501Y potentiation of Q498K (Fig. 2C), which likewise introduces a long aliphatic side chain with a positively charged head group. The complex basis for Q498R/N501Y epistasis (Fig. 4C,D) and the related lack of correlation between epistatic shift and structural displacement suggests that this region’s mutational variability reflects a dynamic basis of ACE2 interaction.

Overall, SARS-CoV-2 has explored a diverse set of mutations during its evolution in humans. Our results show how this ongoing evolution is itself shaping potential future routes of change by shifting the effects of key mutations. Other human coronaviruses have proven adept at escaping from antibody immunity (*25*) because they can undergo extensive evolutionary remodeling of the amino-acid sequence of their receptor-binding domain while retaining high receptor affinity (*26*, *27*). Our work provides large-scale sequence-function maps that help understand how a similar process may play out for SARS-CoV-2.

## Acknowledgements

We thank the Genomics and Flow Cytometry core facilities at Fred Hutchinson Cancer Research Center, Katy Munson at the University of Washington PacBio Sequencing Services, and the Fred Hutch Scientific Computing group supported by ORIP grant S10OD028685. We thank Rachel Eguia for experimental assistance. We thank Steven Weber and the library synthesis team at Twist Bioscience for library construction. We thank Patrick Hernandez and Nadine Czudnochowski for support with protein production, Jay C. Nix for X-ray data collection and Tristan I. Croll for help with structure refinement. Use of the Stanford Synchrotron Radiation Lightsource, SLAC National Accelerator Laboratory, is supported by the DOE, Office of Science, Office of Basic Energy Sciences under Contract No. DE-AC02-76SF00515. The SSRL Structural Molecular Biology Program is supported by the DOE Office of Biological and Environmental Research, and by the NIH NIGMS (including P41GM103393). The contents of this publication are solely the responsibility of the authors and do not necessarily represent the official views of NIGMS or NIH. **Funding:** This project has been funded in part with federal funds from the NIAID/NIH under Contracts No. 75N93021C00015 and HHSN272201400006C, and R01AI1417097 (to JDB); and DP1AI158186 and HHSN272201700059C to DV. Funding was also provided by a Pew Biomedical Scholars Award (DV), an Investigators in the Pathogenesis of Infectious Disease Awards from the Burroughs Wellcome Fund (DV), Fast Grants (DV), and the Natural Sciences and Engineering Research Council of Canada (MM). TNS is an HHMI Fellow of the Damon Runyon Cancer Research Foundation. DV and JDB are Investigators of the Howard Hughes Medical Institute. **Author contributions:** TNS, AJG, and JDB designed the study. TNS and AJG performed deep mutational scanning experiments. TNS analyzed epistasis in the deep mutational scanning data. WWH created interactive data visualizations. TNS and ANL performed pseudotyped lentiviral entry and neutralization experiments, with reagents and assistance from AGF, BD, KAM, and DC. TNS and WWH analyzed the SARS-CoV-2 evolutionary data. EF, JRD, MM, DV, and GS determined the Beta X-ray crystal structure. KH and GS performed and analyzed molecular dynamics simulation. TNS, AJG, and JDB wrote the initial draft, and all authors edited the final version. **Competing Interests:** JDB consults for Moderna on viral evolution and epidemiology and Flagship Labs 77 on deep mutational scanning. TNS, AJG, ANL, and JDB may receive a share of IP revenue as inventors on Fred Hutch-optioned technology/patents related to deep mutational scanning of viral proteins. KH, EF, JRD, DC, and GS are employees of Vir Biotechnology and may hold shares in Vir Biotechnology. **Data and materials availability:** Raw sequencing data are on the NCBI SRA under BioProject PRJNA770094, BioSamples SAMN25941479 and SAMN22208699 (PacBio sequencing) and SAMN25944367 (Illumina barcode sequencing). All code and data at various stages of processing is available together with summaries notebooks detailing the computational pipeline at https://github.com/jbloomlab/SARS-CoV-2-RBD_DMS_variants. Final mutant deep mutational scanning phenotypes are available on GitHub (https://github.com/jbloomlab/SARS-CoV-2-RBD_DMS_variants/blob/main/results/final_variant_scores/final_variant_scores.csv) and Data S1, and interactive visualizations of key data are available at https://jbloomlab.github.io/SARS-CoV-2-RBD_DMS_variants/.

## Materials and Methods

### Library generation

Site-saturation mutagenesis libraries spanning all 201 positions in variant RBDs were created as described for the Beta library by Greaney et al. (*14*). Briefly, a plasmid encoding the Wuhan-Hu-1 RBD (Genbank MN908947, residues N331–T531) was mutagenized to create variant RBDs Alpha (N501Y), Beta (K417N+E484K+N501Y, described in (*14*)), Delta (L452R+T478K), and Eta (E484K). Site saturation mutant libraries in each variant background were produced by Twist Bioscience, introducing precise codon mutations to encode the 20 possible amino acids with no stop codons at each RBD position. Libraries were provided as dsDNA oligos, to which we appended N16 barcodes via PCR primer addition. Failed positions in the Twist mutagenesis were mutagenized in-house using NNS degenerate primers and pooled with the respective Twist libraries for N16 barcoding. Barcoded libraries were cloned into a recipient vector backbone for yeast-surface display (*2*) via Gibson Assembly. Libraries were electroporated into *E. coli* (NEB C3020K) and plated in duplicate at a target bottleneck of 50,000 unique barcodes per library (100,000 for Delta libraries, which were processed subsequent to the other libraries). Colonies were scraped from transformations of each RBD variant library, and plasmid was purified and transformed into the AWY101 yeast strain (*28*).

As described (*2*, *14*), plasmid samples were sequenced using a PacBio Sequel IIe to generate long sequence reads that span the N16 barcode and RBD coding sequence. Raw CCS reads are available on the NCBI Sequence Read Archive, BioProject PRJNA770094, BioSample SAMN25941479 (together with prior Beta PacBio reads under BioSample SAMN22208699 (*14*)). Reads were processed using alignparse (version 0.2.4) (*29*) to generate a table linking each N16 barcode to its unique RBD mutant. Barcode lookup tables for each variant background are available at https://github.com/jbloomlab/SARS-CoV-2-RBD_DMS_variants/tree/main/results/variants and https://github.com/jbloomlab/SARS-CoV-2-RBD_Delta/blob/main/results/variants/codon_variant_table.csv.

### ACE2-binding titration deep mutational scanning experiments

The ACE2-binding affinity of each RBD mutant was determined via massively parallel yeast-surface display titrations (*2*, *30*). Titrations were performed on pooled mutant libraries of the Wuhan-Hu-1, Alpha, Beta, and Eta variants, while the Delta libraries were assayed independently. For the Wuhan-Hu-1, Alpha, Beta, and Eta pools, two experimental replicate titrations were performed on the pooled “replicate 1” libraries, and a single titration was performed on the pooled “replicate 2” libraries. For the Delta library, a single experimental replicate was performed with each library replicate. Frozen yeast libraries were thawed, grown overnight in SD-CAA media (6.7 g/L Yeast Nitrogen Base, 5.0 g/L Casamino acids, 2.13 g/L MES, and 2% w/v dextrose), and backdiluted to 0.67 OD600 in SG-CAA+0.1%D (SD-CAA with 2% galactose + 0.1% dextrose replacing the 2% dextrose) to induce RBD surface expression, which proceeded for 16-18 hours at room temperature with mild agitation.

Induced cells were washed with PBS-BSA (0.2 mg/L), and split into incubations with biotinylated monomeric human ACE2 protein (ACROBiosystems AC2-H82E8) across a concentration range from 10^−6^ to 10^−13^ M at 1-log intervals, plus a 0 M ACE2 sample. Incubations equilibrated overnight at room temperature with mixing. Yeast were washed with ice-cold PBS-BSA and fluorescently labeled with 1:100 FITC-conjugated chicken anti-Myc (Immunology Consultants CMYC-45F) to detect yeast-displayed RBD protein and 1:200 PE-conjugated streptavidin (Thermo Fisher S866) to detect bound ACE2. At each ACE2 sample concentration, single RBD^+^ cells were partitioned into bins of ACE2 binding (PE fluorescence) using a BD FACS Aria II as shown in Fig. S1A. A minimum of 10 million cells were collected at each sample concentration. Cells in each bin were grown overnight in 1 mL SD-CAA+pen-strep, and plasmid was isolated using 96-well yeast miniprep kits (Zymo D2005) according to manufacturer instructions, with the addition of an extended (>2 hr) Zymolyase treatment and a −80°C freeze/thaw cycle prior to cell lysis. N16 barcodes in each post-sort sample were PCR amplified as described in Starr et al. (*2*) and submitted for Illumina HiSeq 50bp single end sequencing. Sequencing reads are available on the NCBI Sequence Read Archive, BioProject PRJNA770094, BioSample SAMN25944367.

Demultiplexed Illumina barcode reads were aligned to library barcodes in barcode-mutant lookup tables using dms_variants (version 0.8.9), yielding a table of counts of each barcode in each FACS bin which is available at https://github.com/jbloomlab/SARS-CoV-2-RBD_DMS_variants/blob/main/results/counts/variant_counts.csv. Read counts in each FACS bin were downweighted by the ratio of total sequence reads from a bin to the number of cells that were sorted into that bin from the FACS log.

We estimated the level of ACE2 binding of each barcoded mutant at each ACE2 sample concentration based on its distribution of counts across FACS bins as the simple mean bin (*31*) as described in (*2*). We determined the binding constant *K*_D_ describing the affinity of each barcoded mutant for ACE2 along with free parameters *a* (titration response range) and *b* (titration curve baseline) via nonlinear least-squares regression using the standard non-cooperative Hill equation relating the mean sort bin to the ACE2 labeling concentration:

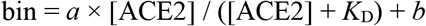

The measured mean bin value at a given ACE2 concentration was excluded from curve fitting for a barcode if fewer than 2 counts were observed across the four FACS bins at that ACE2 concentration. Individual concentration points were also excluded from the curve fit if they exhibited bimodality (>40% of counts of a barcode were found in two non-consecutive bins). To avoid errant fits, we constrained the baseline fit parameter *b* to be between 1 and 1.5, the response parameter *a* between 2 and 3, and the *K*_D_ parameter between 1e-15 and 1e-5. The fit for a barcoded mutant was discarded if the average count across all sample concentrations was below 2, or if more than one sample concentration was missing or excluded. We also discarded curve fits where the normalized mean square residual (residuals normalized relative to the fit response parameter *a*) was >40 times the median normalized mean square residual across all titration fits. Final *K*_D_ binding constants were expressed as −log_10_(*K*_D_), where higher values indicate higher binding affinity. The computational pipeline for computing per-barcode binding constants is available at https://github.com/jbloomlab/SARS-CoV-2-RBD_DMS_variants/blob/main/results/summary/compute_binding_Kd.md.

Because most mutants in the library were independently associated with more than one N16 barcode, we were able to average across internal replicates to derive final mutant affinities. Barcode affinities associated with identical RBD genotypes were first averaged within each experimental replicate. The correlations in per-replicate collapsed affinities are shown in Fig. S1C. The final average was then determined as the average of each replicate value. The final collapsed *K*_D_ for each mutant is given in Data S1 and https://github.com/jbloomlab/SARS-CoV-2-RBD_DMS_variants/blob/main/results/final_variant_scores/final_variant_scores.csv. The median mutant’s ACE2 binding affinity measurement collapses 15 total barcodes across the replicate titration experiments (Fig. S1E). The number of independent barcodes collapsed into each mutation’s final *K*_D_ is reported in the raw data table linked above.

### RBD expression deep mutational scanning experiments

Libraries were grown and induced for RBD expression as described above. Induced cells were washed, labeled with 1:100 FITC-conjugated chicken anti-Myc, and washed in preparation for FACS. Single cells were partitioned into bins of RBD expression (FITC fluorescence) using a BD FACS Aria II as shown in Fig. S1B. A total of >10 million viable cells (estimated by plating of post-sort dilutions) were collected for each library. Cells in each bin were grown out, plasmid isolated, and N16 barcodes sequenced as described above, except read count downweighting used the post-sort colony counts instead of the FACS log counts. Sequencing reads are available on the NCBI Sequence Read Archive, BioProject PRJNA770094, BioSample SAMN25944367.

We estimated the level of RBD expression (mean fluorescence intensity, MFI) of each barcoded mutant based on its distribution of counts across FACS bins and the known log-transformed fluorescence boundaries of each sort bin using a maximum likelihood approach (*2*, *31*), implemented using the fitdistrplus package (version 1.0.14) in R (*32*). Expression measurements were discarded for barcodes for which fewer than 10 counts were observed across the four FACS bins. The full pipeline for computing per-barcode expression values is available at https://github.com/jbloomlab/SARS-CoV-2-RBD_DMS_variants/blob/main/results/summary/compute_expression_meanF.md. Final mutant expression values were collapsed within and across replicates as described above (correlation between replicates shown in Fig. S1D), with a median of 10 barcodes collapsed into each mutant expression measurement (Fig. S1E). Final mutant expression values are available in Data S1 and https://github.com/jbloomlab/SARS-CoV-2-RBD_DMS_variants/blob/main/results/final_variant_scores/final_variant_scores.csv.

### Quantification of epistasis

Epistatic shifts at each site between pairs of RBD variants were quantified via a summary metric, Jensen-Shannon divergence, that captures variation in the 20 amino-acid-level affinity or expression phenotypes measured at a position (*33*). First, affinity (dissociation constant *K*_D_) or expression (MFI) phenotypes *f*_*i*_ of each mutant *i* at a site were transformed to a probability analog *πi*:

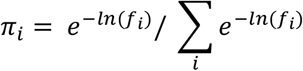

Note that for affinity, the resulting *π*_*i*_ values correspond to Boltzmann weights, since −ln(*K*_D_) is proportional to the free energy in units of *k*_B_*T*. The Jensen-Shannon divergence between vectors *p* and *q* representing the 20 amino acid *π*_*i*_ at a site in two RBD variants is the average Kullback-Leibler divergence of *p* and *q* from their per-index-mean vector *m*:

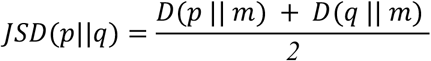

where the Kullback-Leibler divergence is calculated as:

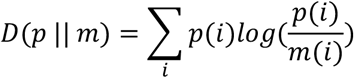

The Jensen-Shannon divergence ranges from 0 for two vectors of probabilities that are identical to 1 for two vectors that are completely dissimilar. For context, scatterplots in Fig. 2C and Figs. S3, S4C illustrate the amino-acid-level epistatic perturbations that give rise to a range of Jensen-Shannon divergence values. To avoid noisier measurements artifactually inflating the epistatic shift metric, a given amino acid mutation was only included in the comparison between a pair of RBD variants if its final measurement was averaged across 3 or more individual barcodes in each RBD background. The calculation of epistatic shifts can be found at https://github.com/jbloomlab/SARS-CoV-2-RBD_DMS_variants/blob/main/results/summary/epistatic_shifts.md.

For epistatic interactions between particular mutations (e.g., Fig. 3A,B,D and Fig. S5D,E), the expected difference in binding of a multiply mutated genotype compared to wildtype in the absence of epistasis is the sum of the Δlog_10_(*K*_D_) measurements of the component single mutations. Any deviation from this addition is a reflection of epistasis. An observed phenotype that is higher than the expected additive phenotype is considered “positive” epistasis, while an observed phenotype that is lower than the additive expectation is considered “negative” epistasis.

### Analysis of SARS-CoV-2 genetic variation

A real-time phylogenetic tree of global SARS-CoV-2 genomes was used to count substitution accrual on internal branches of the SARS-CoV-2 phylogeny. The SARS-CoV-2 mutation-annotated tree described by McBroome et al. (*20*) was downloaded on Feb. 7, 2022. Nucleotide mutations annotated on the tree were converted to amino acid mutations using matUtils (version 0.4.8) (*20*), using the Wuhan-Hu-1 genome (NCBI RefSeq NC_045512.2) as a reference. Amino acid substitutions were tabulated, excluding substitutions that occurred to isolated terminal branches. Substitutions were counted as occurring on N501 versus Y501 genomes, and substitutions that occurred coincidental with a 501 substitution were counted as accruing on the derived 501 state from that branch. A pseudocount of 1 was added to all substitution counts to enable log-ratio comparison of mutation accrual on N501 versus Y501 genomes (Fig. 3E).

### Pseudovirus generation, titering, and neutralization experiments

A spike-pseudotyped lentiviral platform (*34*) was used to measure entry efficiency and antibody neutralization of spike variants. Single and double amino acid mutations and the entire suite of mutations found in Omicron BA.1 (*8*) were introduced into a previously described spike-expression plasmid containing D614G and a 21-amino-acid deletion in the cytoplasmic tail (Addgene 158762) (*34*). Plasmids were purified in triplicate for independent viral rescues from separate plasmid stocks.

Spike pseudotyped lentiviral particles were produced in HEK-293T cells (ATCC CRL-3216). 5e5 cells per well were seeded in 6-well plates in 2mL D10 growth media (Dulbecco’s Modified Eagle Medium with 10% heat-inactivated fetal bovine serum, 2 mM L-glutamine, 100 U/mL penicillin, and 100 μg/mL streptomycin) at 37℃ in a humidified 5% CO_2_ incubator. 24 hours later, cells were transfected using BioT (Bioland Scientific) with 340 ng of spike expression plasmid (or no spike control), 1,135 ng of a Luciferase_IRES_ZsGreen lentiviral backbone, and 865 ng of Gag/Pol lentiviral helper plasmid (BEI Resources NR-52517). Media was changed 24 hours post-transfection. Approximately 65 hours post-transfection, viral supernatants were collected, filtered through a 0.45 μm syringe filter, and stored at −80°C. Particle concentration of each viral supernatant was determined in technical duplicate by p24 ELISA (Advanced Bioscience Laboratories Cat. # 5421) versus a known particle standard according to manufacturer instructions.

Entry titers of spike-pseudotyped lentiviral particles were determined on HEK-293T cell lines expressing high levels of human ACE2 (BEI Resources NR-52511) (*34*) and low levels of human ACE2. The low level ACE2 cells were created by genomically integrating single AttB_ACE2-miRFP670_IRES_mCherry-H2A-P2A-PuroR plasmids into HEK-293T LLP-Int-BFP-IRES-iCasp9-Blast Bxb1 landing pad cells (*35*), followed by selection with 1 μg/mL puromycin. Low ACE2 levels were achieved by using a “AATTTT[ATG]” Kozak sequence preceding ACE2 to decrease its steady-state abundance, unmasking entry deficits (*35*). All cells were grown in D10 media, and the ACE2-low cell line was supplemented with 1 μg/mL doxycycline to maintain ACE2 expression. To quantify ACE2 expression levels, HEK-293T cells expressing high, low or no ACE2 were resuspended in FACS buffer (PBS+2% BSA) and incubated for 1 h with 1:500 rabbit anti-ACE2 antibody (Abcam ab272500). Cells were washed with FACS buffer and labeled for 1 h with 1:3000 Alexa-Fluor-488 goat anti-rabbit IgG H&L (Abcam ab150077). Cells were washed with FACS buffer and fixed with 4% PFA. ACE2 expression (AF488 fluorescence) was measured via flow cytometry on a BD LSRFortesssa X50, and geometric mean fluorescence intensity was compared between samples (Fig. S6A).

For titering, ACE2-high and ACE2-low cells were seeded at 1.2e4 cells per well in poly-L-lysine-coated 96-well plates (Greiner 655930) in 50 uL D10 media (plus 1 μg/mL doxycycline for the ACE2-low cell line). 24 hours later, 100 μL of viral supernatants prepared across a 2-fold dilution series (1:2 to 1:256) were added to cells. Approximately 65 hours post-infection, luciferase activity was measured (Promega Bright-Glo, E2620). Relative luciferase unit (RLU) measures were averaged across viral dilutions within linear range and expressed relative to p24 particle concentration as the final p24-normalized entry titer (RLU/pg p24).

Antibody neutralization of spike-pseudotyped particles was measured on ACE2-high cell lines in technical duplicate. Cells were seeded in 96-well plates as described above for virus titering. 24 hours later, viral supernatants were diluted to a to a target of 400,000 RLU per well, and incubated for 1 hour at 37°C with monoclonal antibody across 8 four-fold dilution points starting at a concentration 400-fold higher than the expected IC50 (*16*). 100 μL of antibody-virus mixture was added to cells, and luciferase activity was measured ~65 hours post-infection as described for titering experiments. Fraction infectivity of each sample was calculated relative to a no-antibody well inoculated with the same viral supernatant in a matched row of the 96-well plate. We used neutcurve (https://jbloomlab.github.io/neutcurve, version 0.5.7) to calculate the inhibitory concentration 50% (IC50) of each antibody against each virus by fitting a Hill curve with fixed baseline 0 and plateau 1.

### Recombinant protein production

SARS-CoV-2 Beta RBD for crystallization (residues 328-531 of S protein from GenBank NC_045512.2 with N-terminal signal peptide and C-terminal 8xHis-tag) was expressed in Expi293F (Thermo Fisher Scientific) cells in the presence of 10 µM kifunensine at 37°C and 8% CO2. Transfection was performed using the ExpiFectamine 293 Transfection Kit (Thermo Fisher Scientific). Cell culture supernatant was collected four days after transfection and supplemented with 10x PBS to a final concentration of 2.5x PBS (342.5 mM NaCl, 6.75 mM KCl and 29.75 mM phosphates). SARS-CoV-2 Beta RBD was purified using a 5 mL HisTalon Superflow cartridge (Takara Bio) followed by buffer exchange into PBS using a HiPrep 26/10 desalting column (Cytiva).

Human ACE2 for crystallization (residues 19-615 from Uniprot Q9BYF1 with a C-terminal thrombin cleavage site-TwinStrep-10xHis-GGG-tag, and N-terminal signal peptide) was expressed in Expi293F cells in the presence of 10 µM kifunensine at 37°C and 8% CO2. Transfection was performed using the Expi293 transfection kit (Thermo Fisher Scientific). Cell culture supernatant was collected five days after transfection and supplemented to a final concentration of 80 mM Tris-HCl pH 8.0, 100 mM NaCl, and then incubated with BioLock (IBA GmbH) solution. ACE2 was purified using a 1 mL StrepTrap HP column (Cytiva). Protein-containing fractions were pooled and digested with EndoH and thrombin at 4°C overnight. ACE2 was further purified by size exclusion chromatography using a Superdex 200 Increase 10/300 GL column (Cytiva) equilibrated in 20 mM Tris-HCl pH 7.5, 150 mM NaCl.

### Crystallization, data collection, structure determination, and analysis

SARS-CoV-2 Beta RBD was mixed with a 1.4-fold molar excess hACE2, and a 1.3-fold molar excess of S304 Fab and S309 Fab. The complex was purified on a Superdex 200 10/300 GL column pre-equilibrated in 20 mM Tris-HCl pH 7.5, 150 mM NaCl. Crystals of the SARS-CoV-2 Beta RBD-hACE2-S304-S309 Fab complex were obtained at 20°C by sitting drop vapor diffusion. A total of 200 nL of the complex at 6 mg/mL was mixed with 200 nL mother liquor solution containing 10% w/v PEG 8000, 20% v/v ethylene glycol, 0.1 M Tris (base)/bicine pH 8.5, 3% w/v D-sorbitol. Crystals were flash frozen in liquid nitrogen.

Data were collected at Beamline 9-2 of the Stanford Synchrotron Radiation Lightsource facility in Stanford, CA and processed with the XDS software package (*36*) yielding a final dataset of 2.45 Å in space group P1. The SARS-CoV-2 Beta RBD-hACE2-S304-S309 complex structure was solved by molecular replacement using Phaser (*37*) from a starting model consisting of ACE2-RBD-S304-S309 (PDB: 7L0N). Several subsequent rounds of model building and refinement were performed using Coot (*38*), ISOLDE (*39*), Refmac5 (*40*), Phenix (*41*) and MOE (https://www.chemcomp.com), to arrive at a final model of the quaternary complex.

### MD simulation

Coordinates of the Wuhan-Hu-1 RBD:hACE2 structure were prepared as previously described (*42*), where the RBD was taken from PDB 6M0J, hACE2 from PDB 1R42, and their complex generated by aligning the proteins to the CST 6M0J structure. Complex glycans were then added to the structure at positions 53, 90, 103, 322, 432, 546, and 690 on hACE2 and 343 on the RBD followed by refinement with ISOLDE (*39*). These coordinates were then prepared using QuickPrep (MOE v2020.0901, https://www.chemcomp.com). The Omicron RBD:hACE2:S304:S309 structure from PDB 7TN0 was glycosylated as described above and prepared using QuickPrep.

The coordinates of each RBD:ACE2 complex were parameterized using tleap (*43*). The following force fields were used: Amber ff14SB for the protein (*44*), GLYCAM_06j-1 for glycans (*45*), TIP3P for water (*46*) and for the neutralizing 0.15 M of NaCl Joung & Cheatham parameters (*47*) were used, and the Li parameters (*48*) for Zn and divalent ions.

For each complex, a nine-stage restrained minimization and equilibration protocol was used as previously described (*49*) using Amber20 (*43*). Initial minimization involved 10,000 steps and positional restraints applied to all heavy atoms resolved in the crystal structures with a 100 kcal/molÅ^2^ force constant. 8 independent simulations were initialized using different initial velocities drawn randomly from the Maxwell-Boltzmann distribution, in an NVT ensemble, gentle heating from 100 K to 300 K over 100 ps, and positional restraints on all heavy atoms with a 100 kcal/molÅ^2^ force constant. Next the ensemble was switched to NPT, a constant temperature of 300 K over 100 ps, and positional restraints on all heavy atoms with a 100 kcal/molÅ^2^ force constant. Next, a 250 ps stage was run in NPT at 300 K with positional restraints on all heavy atoms with a 10 kcal/molÅ^2^ force constant. Next minimization for 10,000 steps was performed with positional restraints on backbone atoms with a 10 kcal/molÅ^2^ force constant. This was followed by 100 ps of NPT at 300 K with positional restraints on backbone atoms with a 10 kcal/molÅ^2^ force constant. Next 100 ps of NPT at 300 K with positional restraints on backbone atoms with a 1.0 kcal/molÅ^2^ force constant. Next 100 ps of NPT at 300 K with positional restraints on backbone atoms with a 0.1 kcal/molÅ^2^ force constant. The final stage of equilibration was 2.5 ns of unrestrained MD in NPT at 300 K for each of the 8 independent trajectories for each complex. The 8 pairs of simulations were extended for 700 ns each in NPT at 300 K. Snapshots were saved every 1 ns for post-processing. 5.6 μs of aggregated MD simulations were obtained for Omicron RBD:hACE2:S304:309 and 5.6 μs for Wuhan-Hu-1 RBD:hACE2.

Simulations were post-processed using cpptraj (*50*). Coordinates were imaged, RMS-aligned using Cα atoms to the respective crystal structure coordinates, water and ions stripped and saved to netcdf files. Distances were computed using the atoms of Q (OE1, HE1/2, NE2), N (OD1, HD1/2), D (OD1 or OD2), E (OE1 or OE2), R (HE, HH11/2, HH21/2), and K (HZ1/2/3). For each residue pair (H-bond or salt bridge) all distance pairs between atoms were computed and the lowest value of those pairs used as the hydrogen bond or salt bridge distance. Figures were generated using matplotlib.

Volumetric maps were computed using the VolMap Plugin in VMD (*51*) [Humphrey, W. F., Dalke, A. & Schulten, K. (1996) *J. Mol. Graphics* **14,** 33–38.]. Default parameters were used to compute the atomic densities observed over a grid, where the width of gaussian functions centered at each grid point bore widths equal to the atomic radii in each respective residue and then weighted by the atomic mass. The double sum of these gaussians over the grid points and over the course of the simulation were used to generate an isosurface map. The isosurfaces were rendered using an occupancy threshold of 0.5.

### Comparisons among variant RBD structures

We analyzed atomic displacements between structures of ACE2-bound variant RBDs. X-ray crystallography structures included Wuhan-Hu-1: PDB 6M0J (*23*); Alpha: PDB 7EKF (*24*); Beta: PDB 7EKG (*24*); and Delta: PDB 7WBQ (*52*). Cryo-EM structure local refinements included Wuhan-Hu-1: PDB 7KMB (*53*); Alpha: PDB (*54*); Beta: PDB 7VX4 (*55*); and Delta: PDB 7V8B. Structures were aligned in PyMol to minimize RBD structure RMSD. Pairwise distances between Cα or all-atom-averages (for unmutated sites) between aligned structures were computed using the bio3d package in R (*56*).

### Data visualization

Interactive visualizations available at https://jbloomlab.github.io/SARS-CoV-2-RBD_DMS_variants/ were built using altair, version 4.2 (*57*). Sites of strong antibody escape (Fig. 2A and Fig. S4A) were those with an average normalized total escape of >0.05 across all antibodies in the aggregate tool described by (*9*) as of November 27, 2021 (https://raw.githubusercontent.com/jbloomlab/SARS2_RBD_Ab_escape_maps/03910f5bb6bc86ab823e9cf40f34b07b403f26d2/processed_data/escape_data.csv).

## Supplemental Figures

**Fig. S1.**
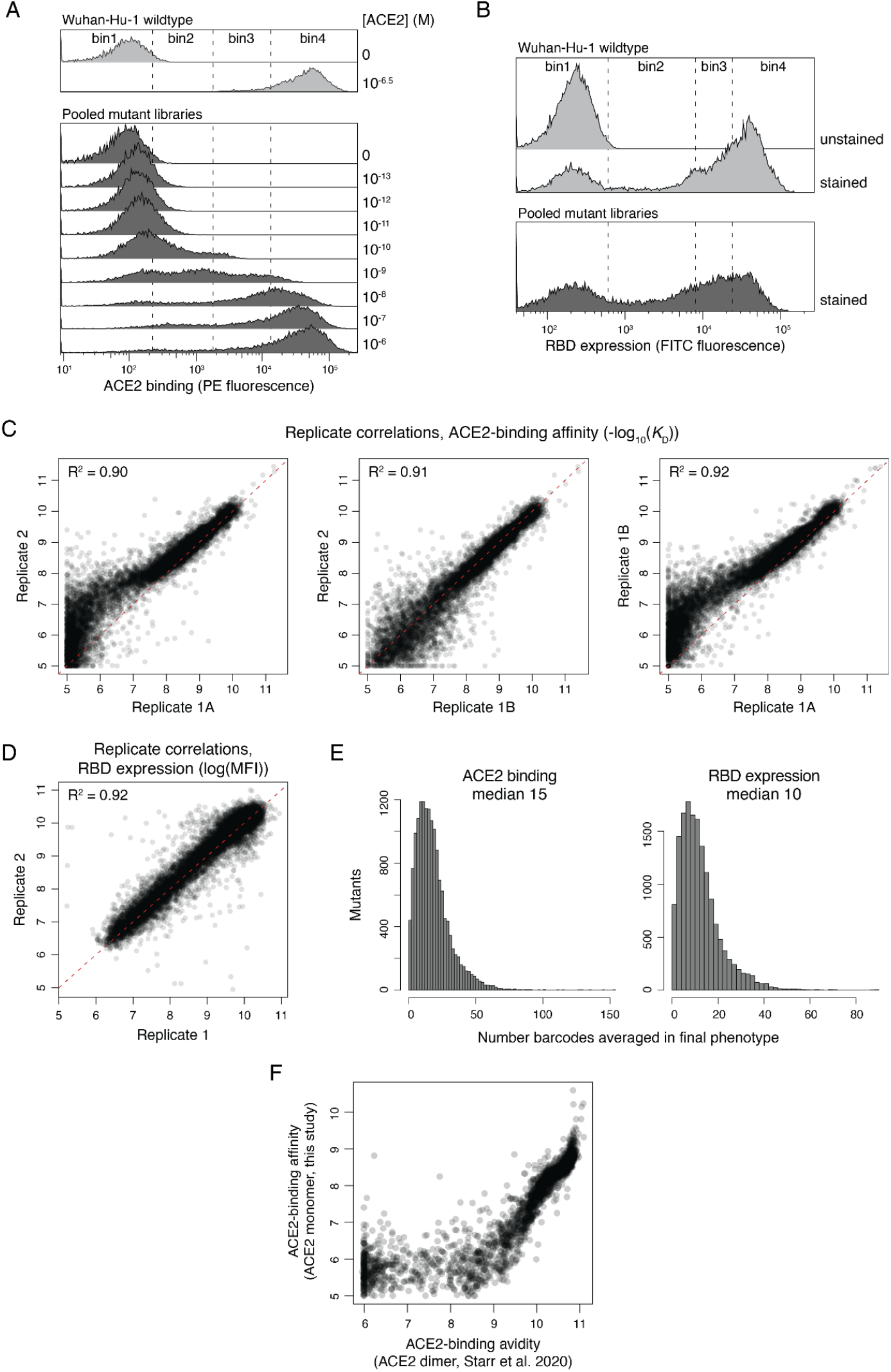
Deep mutational scanning experimental details. (**A**) Representative flow cytometry traces and gates used in ACE2 binding titration FACS-seq assays on the pooled variant libraries. Gates were drawn to isolate single, RBD-positive cells, followed by binning on histograms of ACE2 binding (PE fluorescence) as shown. **(B)** Representative flow cytometry traces and gates used in RBD expression FACS-seq assays on the pooled variant libraries. Gates were drawn to isolate single cells, followed by binning on histograms of RBD expression (FITC fluorescence) as shown. (**C**) Correlations in mutant affinities as measured in triplicate ACE2 titration assays with pooled variant libraries. Red line, 1:1. Replicates 1A and 1B are independent experimental replicates performed on a single library, and replicate 2 is an experiment on an independently generated replicate library. (**D**) Correlations in mutant RBD expression levels as measured in duplicate Sort-seq assays with pooled variant libraries. Red line, 1:1. (**E**). Histograms of the number of barcodes that were averaged for the calculation of final deep mutational scanning scores for each amino acid mutant. (**F**) Relationship between the Wuhan-Hu-1 mutant ACE2-binding affinities as measured in the current study with monomeric ACE2 versus the prior avidity values measured in Starr et al. 2020 using dimeric ACE2 (*2*).

**Fig. S2.**
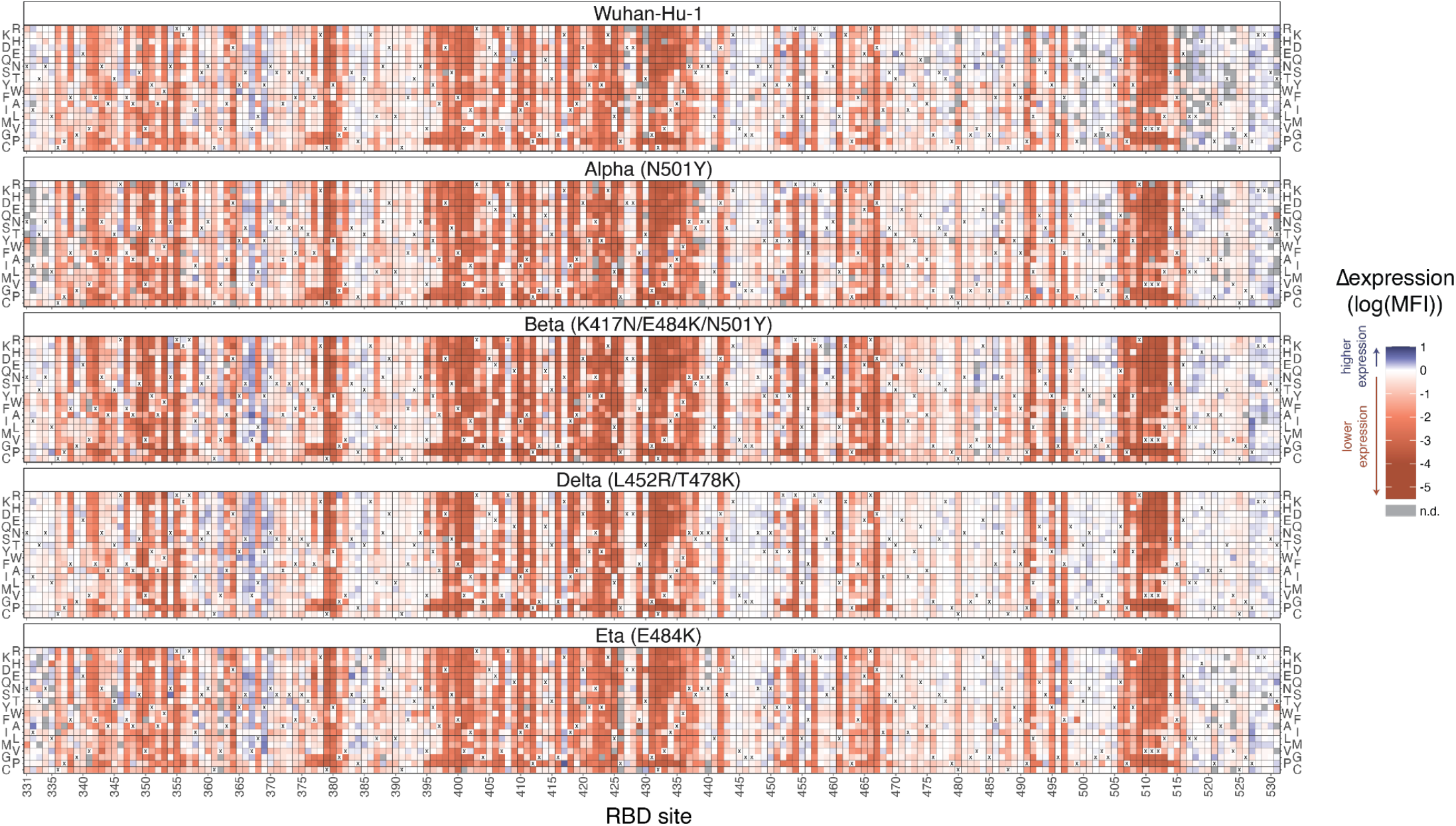
Deep mutational scanning maps of mutational effects on RBD expression. The impact of every single amino-acid mutation in SARS-CoV-2 variant RBDs on yeast surface-expression levels as determined by FACS-seq assays (Fig. S1). The wildtype amino acid in each variant is indicated with an “x”, and gray squares indicate missing mutations in each library. An interactive version of this map is at https://jbloomlab.github.io/SARS-CoV-2-RBD_DMS_variants/RBD-heatmaps/, and raw data are in Data S1.

**Fig. S3.**
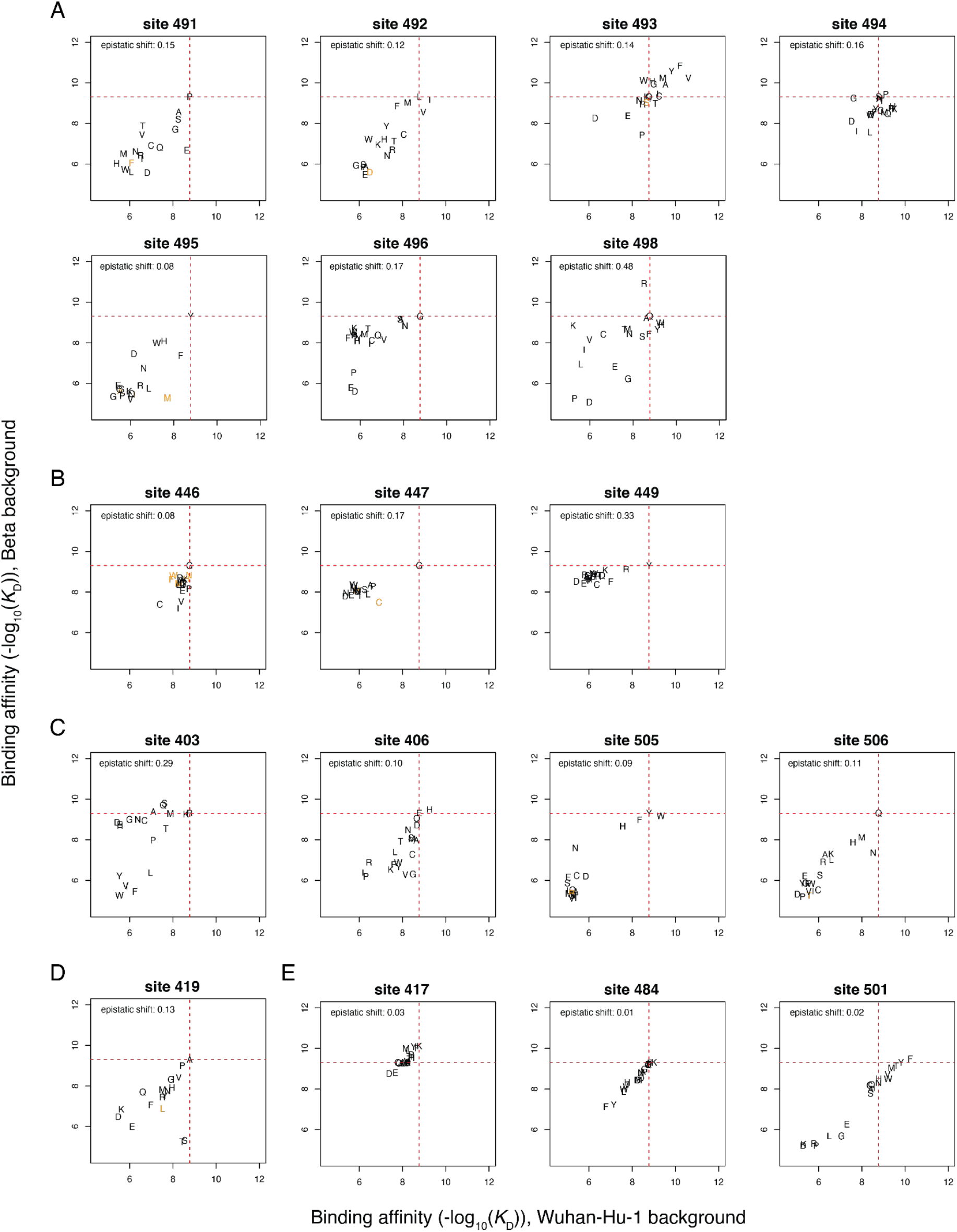
Mutation-level perturbations at sites of interest between the Beta and Wuhan-Hu-1 RBDs. Scatter plots of mutation-level affinities for sites of interest: (**A**) 498 and the central ACE2-contact strand, (**B**) 446-449 loop, (**C**) site 403 and related sites, (**D**) site 419 where mutations to S/T add a glycan in the Beta RBD containing K417N, (**E**) and additive interaction among the sites mutated between Wuhan-Hu-1 and Beta. Details as in Fig. 2C. Orange letters are mutations that were sampled with fewer than 3 unique barcodes across titration replicates in the Beta and/or Wuhan-Hu-1 data and were therefore not included in the epistatic shift computation as described in Methods. Scatterplots for all sites can be visualized at https://jbloomlab.github.io/SARS-CoV-2-RBD_DMS_variants/epistatic-shifts/.

**Fig. S4.**
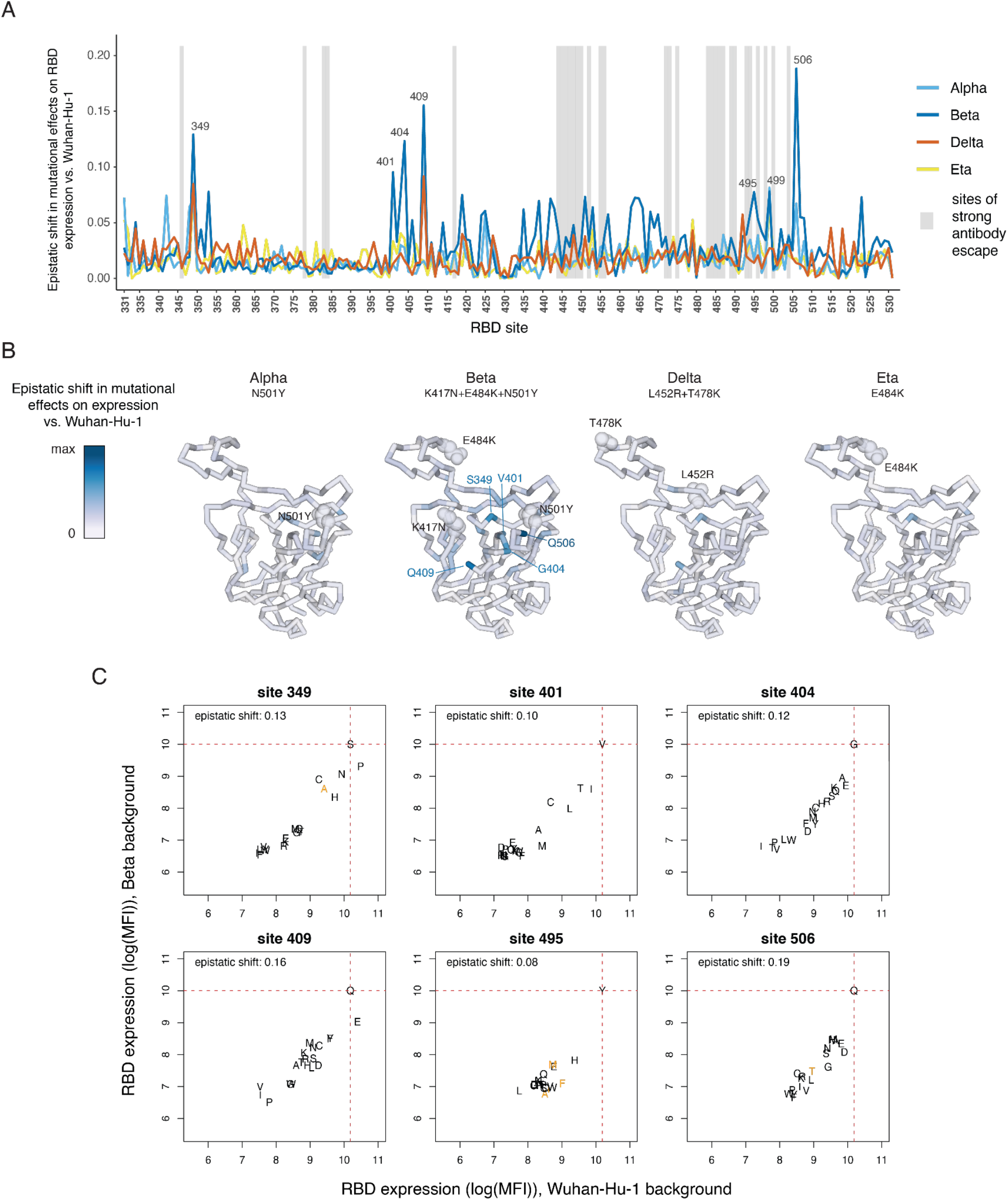
Epistatic shifts in mutational impacts on RBD expression. (**A**) The epistatic shift in mutational effects on RBD expression at each RBD site between the indicated variant and Wuhan-Hu-1. The epistatic shift is calculated as in Fig. 2A (see Methods). (**B**) Structural projection of the epistatic shifts onto the backbone of the Wuhan-Hu-1 RBD structure (PDB 6M0J). Details as in Fig. 2B. (**C**) Scatters of mutation-level RBD expression values for sites of strong epistatic shifts in (B). Details as in Fig. 2C and Fig. S3. These scatter plots suggest that at the shifted sites, the Beta RBD is more sensitive to mutation compared to Wuhan-Hu-1, reflected in the concave shape of each plot. This is in contrast to idiosyncratic epistatic shifts in individual mutation’s effects on ACE2 binding as seen in e.g. Fig. 2C.

**Fig. S5.**
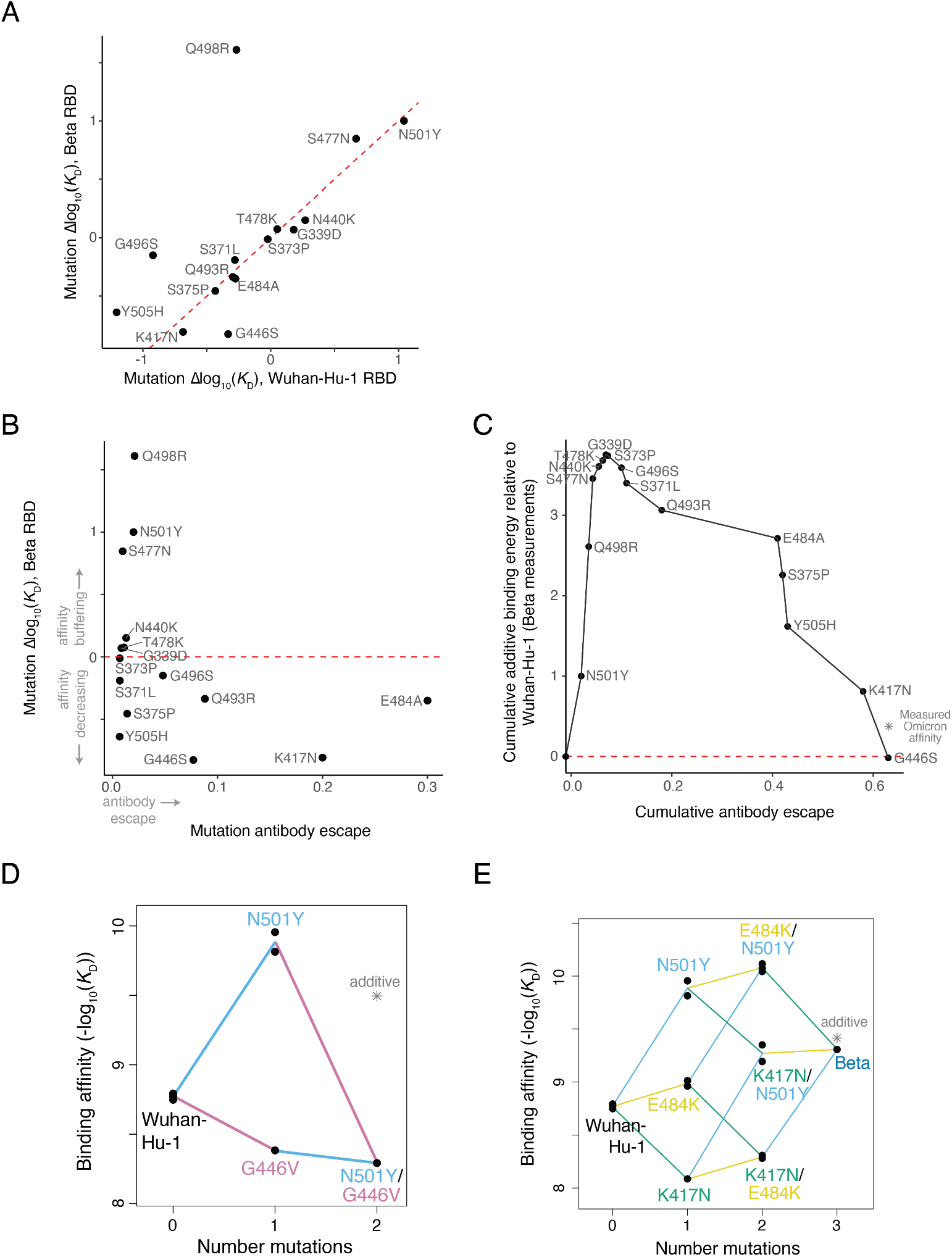
Epistasis and affinity-buffering of Omicron BA.1 RBD mutations. (**A**) Scatter plot illustrating the difference in effect on ACE2 binding for each of the 15 Omicron BA.1 mutations (Fig. 3B) when introduced in the Beta versus Wuhan-Hu-1 RBD. Red line, 1:1. (**B**) Relationship between mutation effects on ACE2 binding versus antibody escape for the fifteen Omicron BA.1 RBD mutations (Fig. 3B). Antibody escape is estimated from a calculator that aggregates >250 antibody deep mutational scanning escape profiles to determine the expected antigenic effect of mutations (*9*). (**C**) Cumulative ACE2-binding affinity versus antibody escape of Omicron BA.1 mutations (Fig. 3B and (B)). Antibody escape calculated as in (B), except mutations were introduced consecutively into the calculator in the order shown, which accounts for possible redundancy in escape mutations at overlapping antibody epitopes. (**D**) Double mutant cycle diagram illustrating negative sign epistasis between N501Y and G446V. (**E**) Triple mutant cycle illustrating additivity among the three mutations in the Beta RBD. See also Fig. S4E.

**Fig. S6.**
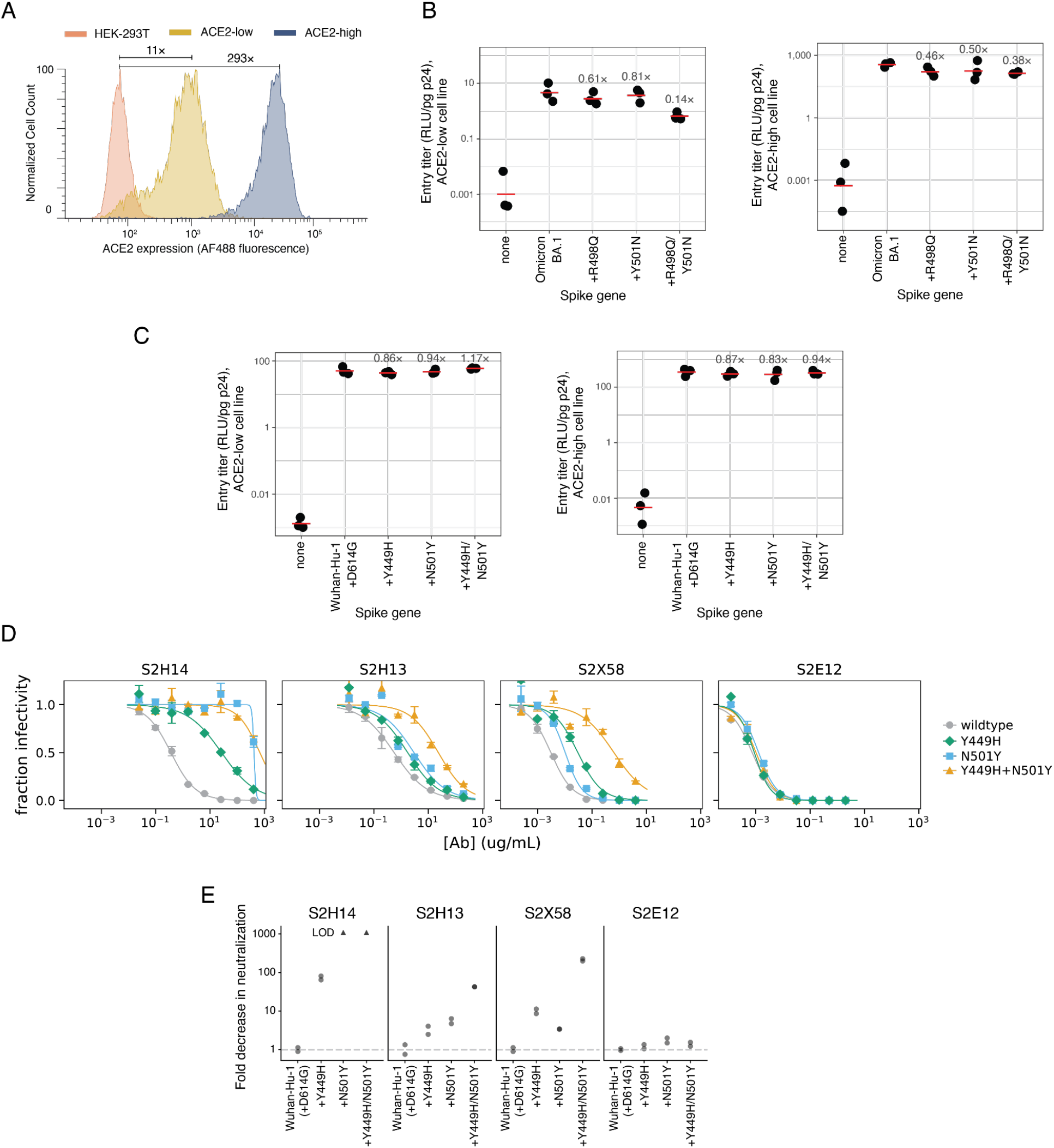
Pseudoviral entry and neutralization assays. (**A**) ACE2 expression of HEK-293T cell lines, determined by antibody labeling of surface ACE2 expression and flow cytometry. Label indicates fold-increase in fluorescence geometric mean. (**B**) Normalized entry titers of Omicron BA.1 (or reversion mutant) spike-pseudotyped lentiviral particles on HEK-293T cell lines expressing low (left; independent titering replicate of Fig. 3C) or high (right) levels of ACE2. Labels indicate fold-change in geometric mean entry (red bar) across biological triplicate measurements. (**C**) Normalized entry titers of Wuhan-Hu-1+D614G (and RBD mutant) spike-pseudotyped lentiviral particles on HEK-293T cell lines expressing low (left) or high (right) levels of ACE2. (**D**) Neutralization of spike-pseudotyped lentiviral particles by human monoclonal antibodies S2H14, S2H13, and S2X58 that showed escape at site Y449 in prior deep mutational scanning data (*16*). Each point is the mean and standard error from technical duplicates. S2E12 is a negative control antibody, which was not expected to be escaped by the Y449H or N501Y mutations based on prior deep mutational scanning data (*16*). (**E**) Fold-change in neutralization of spike-pseudotyped lentivirus from the curves in (D). The Y449H/N501Y double mutant shows 2.4x and 6.3x synergistic escape from S2H13 and S2X58, respectively, compared to the multiplicative fold-change of the single mutants.

**Fig. S7.**
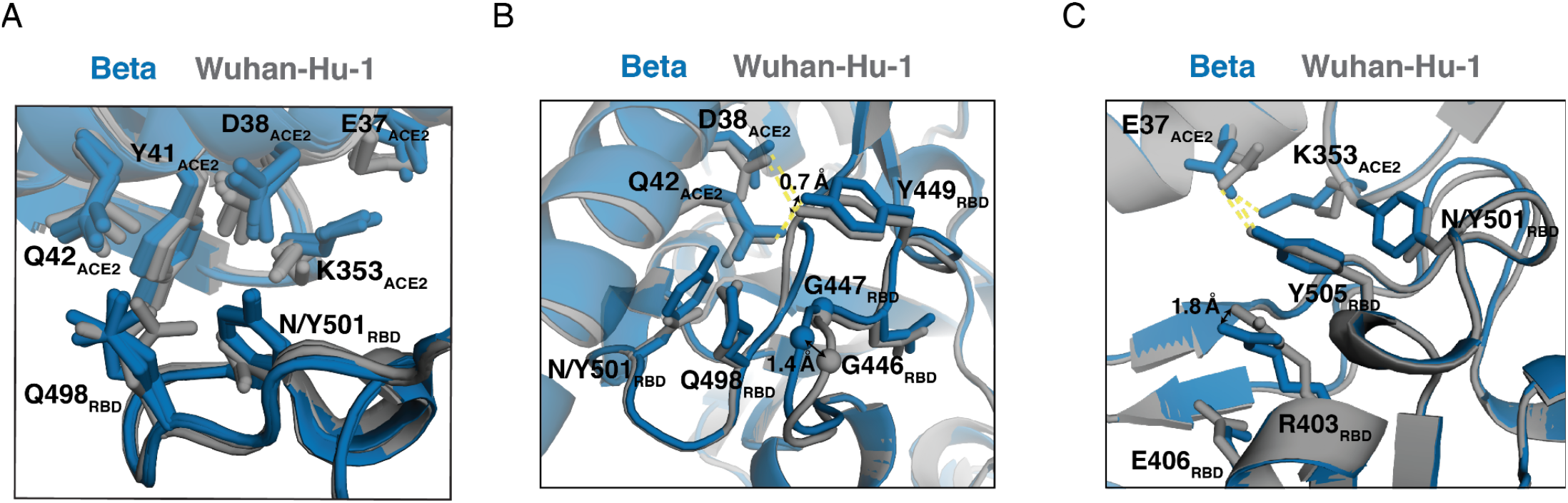
Structural comparison of epistatically shifted sites between Wuhan-Hu-1 and Beta RBDs. Zoomed structural views of clusters of epistatically shifted sites visualized in Wuhan-Hu-1 and Beta ACE2-bound RBD X-ray crystal structures. (**A**) Site 498 shows dramatic epistatic shifts in the presence of N501Y. Comparison of multiple crystal structures of Wuhan-Hu-1 (PDBs 6M0J, 6VW1 [Wuhan-Hu-1 RBM chimera on SARS-CoV-1 RBD scaffold]) and Beta (this study and PDB 7EKG) illustrates overlapping heterogeneity in Q498 rotamers between Wuhan-Hu-1 and Beta structures. (**B**) Sites 446, 447, and 449 exhibit small shifts in backbone and side chain positions in the ACE2-bound structure, though Delta exhibits similar variability in backbone and sidechain positions (see Fig S8B,C) despite lacking epistatic shifts at these positions. (**C**) The epistatically shifted residues 505, 403, and 406 show little structural perturbation despite large shifts in mutational effects, especially at residue 403 (see Fig. 2C). Site 403 has not previously been implicated in key events in the functional or antigenic evolution of SARS-related coronaviruses, but mutations in this region have been shown to transmit allosterically to distal regions of the ACE2-binding surface (*58*, *59*), which may be further reflected in many epistatic shifts in mutation effects on RBD expression in this region (Fig. S4).

**Fig. S8.**
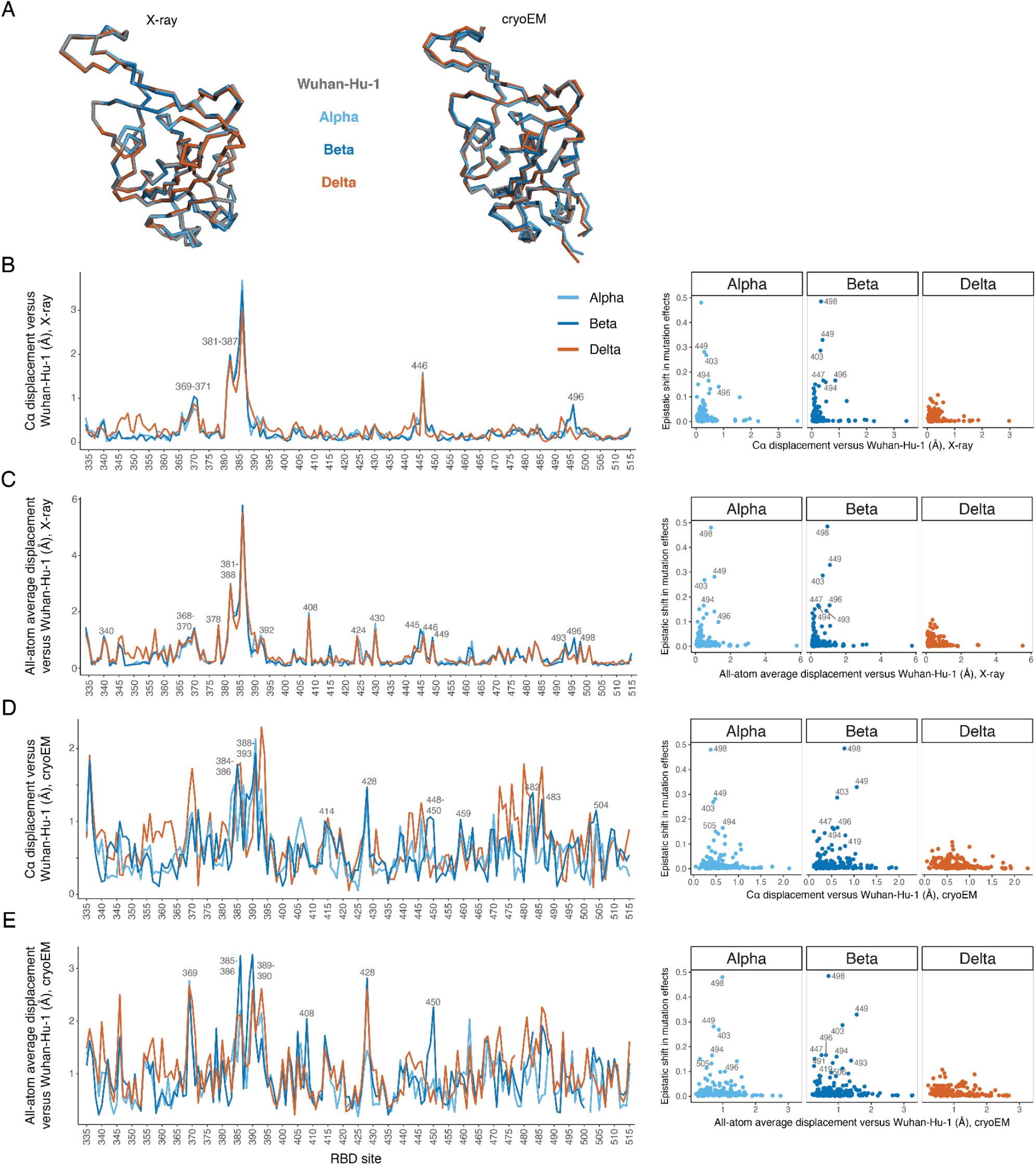
Structural perturbations among variant RBD structures. (**A**) Backbone structure alignment between ACE2-bound RBD X-ray crystal structures (left; Wuhan-Hu-1 (PDB 6M0J, 2.45Å resolution), Alpha (7EKF, 2.85Å), Beta (7EKG, 2.63Å), and Delta (7WBQ, 3.34Å)) or local refinement maps from ACE2-bound spike cryo-EM structures (right; Wuhan-Hu-1 (PDB 7KMB, 3.39Å resolution), Alpha (7MJN, 3.29Å), Beta (7VX4, 3.90Å) and Delta (7V8B, 3.2Å)). (**B**-**E**) Changes in backbone and sidechain properties in variant versus Wuhan-Hu-1 structures. For each property, the left hand plot shows the value for each variant RBD versus Wuhan-Hu-1 as a lineplot across RBD sites; sites with the largest displacements in the Beta structures are labeled. Right Hand plot shows relationship across RBD sites between the property and epistatic shift from deep mutational scanning (Fig. 2A). Calculated properties include the mainchain Cα displacement from X-ray (B) or cryoEM (D) structures, and all-atom average displacement from X-ray (C) or cryoEM (E) structures.

**Fig. S9.**
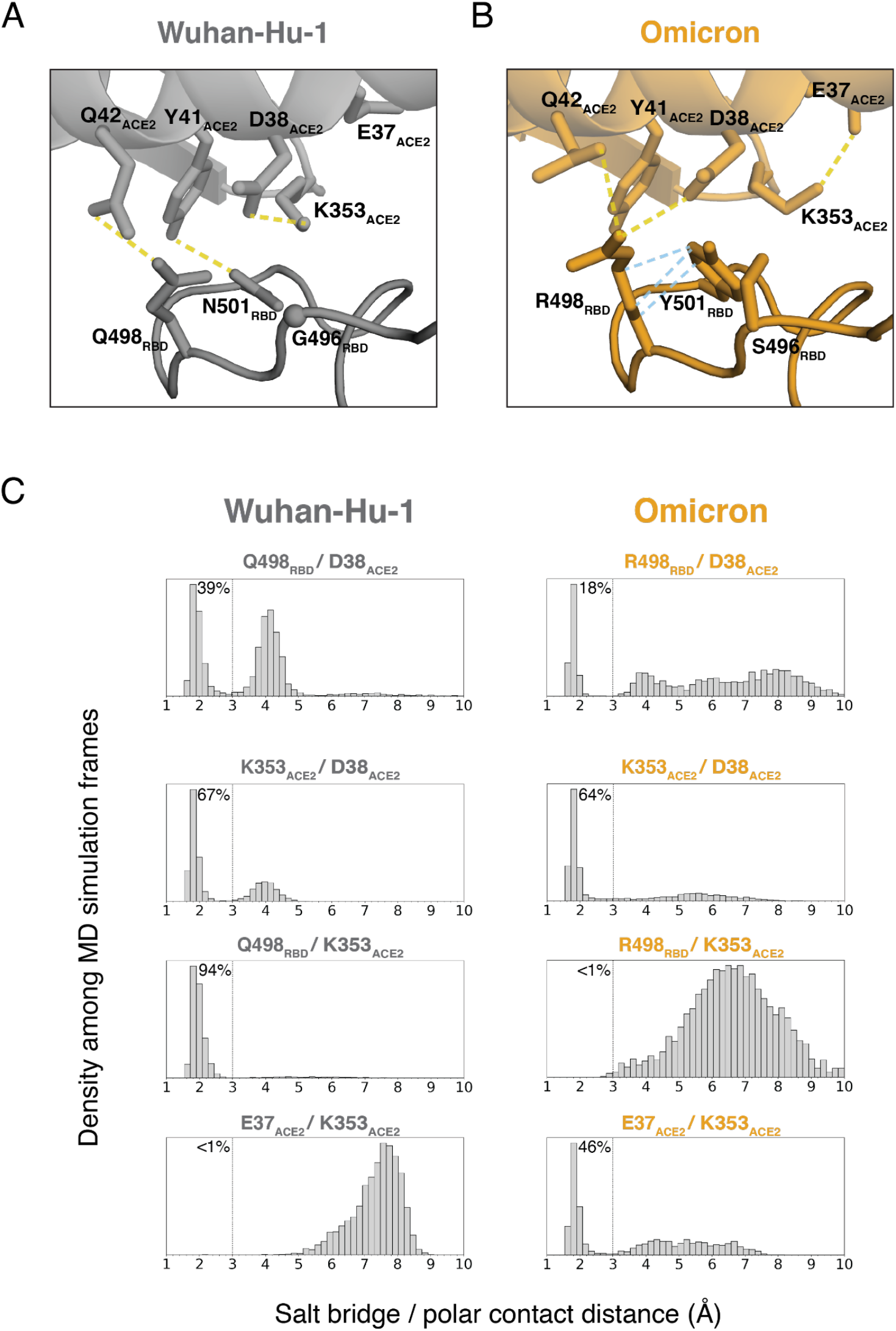
Dynamic RBD:ACE2 interface as revealed in molecular dynamics simulation. (**A**, **B**) Structural context of residues 498 and 501 in the Wuhan-Hu-1 (PDB 6M0J) (C) and Omicron (PDB 7TN0) structures (D). Yellow dash lines indicate polar contacts, and blue dash lines indicate non-polar packing contacts. (**C**) Histograms showing the distances between charged atoms or hydrogen bond donors/acceptors over the course of molecular dynamics simulation of Wuhan-Hu-1 (left) or Omicron (right) RBD bound to ACE2, with the percentage of frames where residues are within 3.0Å distance (dashed line) labeled.

**Table S1.**
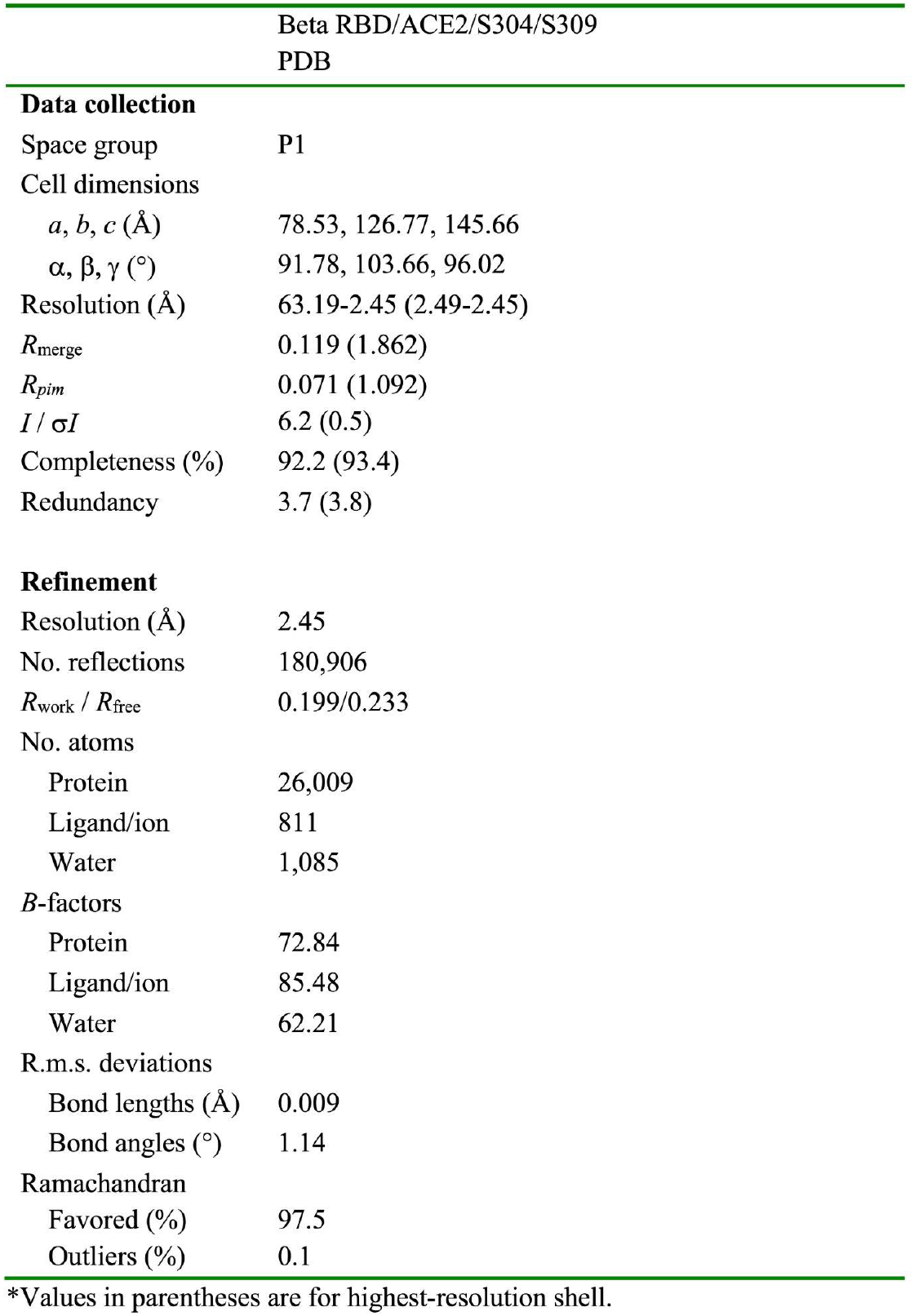
Crystallographic data collection and refinement statistics.

|  | Beta RBD/ACE2/S304/S309<br>PDB |
| --- | --- |
| **Data collection** |  |
| Space group | P1 |
| Cell dimensions |  |
| *a*, *b*, *c* (Å) | 78.53, 126.77, 145.66 |
| $\alpha$, $\beta$, $\gamma$ (°) | 91.78, 103.66, 96.02 |
| Resolution (Å) | 63.19-2.45 (2.49-2.45) |
| *R*<sub>merge</sub> | 0.119 (1.862) |
| *R*<sub>pim</sub> | 0.071 (1.092) |
| *I* / $\sigma I$ | 6.2 (0.5) |
| Completeness (%) | 92.2 (93.4) |
| Redundancy | 3.7 (3.8) |
| **Refinement** |  |
| Resolution (Å) | 2.45 |
| No. reflections | 180,906 |
| *R*<sub>work</sub> / *R*<sub>free</sub> | 0.199/0.233 |
| No. atoms |  |
| Protein | 26,009 |
| Ligand/ion | 811 |
| Water | 1,085 |
| *B*-factors |  |
| Protein | 72.84 |
| Ligand/ion | 85.48 |
| Water | 62.21 |
| R.m.s. deviations |  |
| Bond lengths (Å) | 0.009 |
| Bond angles (°) | 1.14 |
| Ramachandran |  |
| Favored (%) | 97.5 |
| Outliers (%) | 0.1 |

\*Values in parentheses are for highest-resolution shell.

